# FRED: a universal tool to generate FAIR metadata for omics experiments

**DOI:** 10.64898/2026.02.03.703529

**Authors:** Jasmin Walter, Carsten Kuenne, Noah Knoppik, Philipp Goymann, Mario Looso

**Affiliations:** Bioinformatics Core Unit (BCU), Max Planck Institute for Heart and Lung Research, Bad Nauheim, Germany; Cardio-Pulmonary Institute (CPI), Bad Nauheim, Germany

## Abstract

Scientific research relies on transparent dissemination of data and its associated interpretations. This task encompasses accessibility of raw data, its metadata, details concerning experimental design, along with parameters and tools employed for data interpretation. Production and handling of these data represents an ongoing challenge, extending beyond publication into individual facilities, institutes and research groups, often termed Research Data Management (RDM). It is foundational to scientific discovery and innovation, and can be paraphrased as Findability, Accessibility, Interoperability and Reusability (FAIR). Although the majority of peer-reviewed journals require the deposition of raw data in public repositories in alignment with FAIR principles, metadata frequently lacks full standardization. This critical gap in data management practices hinders effective utilization of research findings and complicates sharing of scientific knowledge. Here we present a flexible design of a machine-readable metadata format to store experimental metadata, along with an implementation of a generalized tool named FRED. It enables i) dialog based creation of metadata files, ii) structured semantic validation, iii) logical search, iv) an external programming interface (API), and v) a standalone web-front end. The tool is intended to be used by non-computational scientists as well as specialized facilities, and can be seamlessly integrated in existing RDM infrastructure.

## Introduction

Research Data Management (RDM) constitutes a fundamental component of contemporary scientific practice, encompassing the entire lifecycle of data, including sample collection, data generation, organization, storage, access, and dissemination, along with the management of associated metadata. Due to substantial financial and resource expenditures associated with the generation of omics data, maximizing the utility of these datasets through reuse has emerged as a critical objective, thereby driving growing interest in the integration and repurposing of data from public repositories^1^. However, without proper RDM, experimental designs may be flawed, errors—both random and systematic—can go undetected, and results may be misinterpreted, undermining reliability. In contrast, robust RDM enhances data interpretability and supports independent validation, increasing confidence in research outcomes. Comprehensive metadata furnish the essential contextual information required for the accurate interpretation of data, such as sample origin, experimental conditions, and data processing procedures. It is furthermore mandatory for long-term preservation of scientific knowledge, mitigating the risk of information loss associated with personnel turnover or institutional transitions^2,3^.

The growing emphasis on data-driven science has further heightened the demand for standardized, FAIR (Findable, Accessible, Interoperable, Reusable) data practices. Introduced in 2016 by a coalition of industry, academia, and funding agencies, the FAIR principles aim to ensure that scientific data is not only accessible but also semantically rich, machine-readable, and reusable across disciplines and platforms^4^. The dynamic nature of experimental workflows—spanning diverse omics technologies, hierarchical designs, and evolving methodologies—demands a flexible framework that can adapt to new data types and requirements without compromising consistency or searchability. Illustrative examples for established standards are the Minimum Information About a Microarray Experiment (MIAME)^3^, Minimum Information About a High-throughput Sequencing Experiment (MINSEQE)^5^, and Minimum Information About a Proteomics Experiment (MIAPE)^2^. Of note, a subset of these standards was adopted by relevant trade journals, which subsequently mandated the documentation of high-throughput data in accordance with these standards. This precipitated the implementation of tools like MARMoSET^6^ for the creation of standard-compliant reports, which concentrate on the technical documentation of a measurement, while neglecting to consider the experimental design. Other tools were introduced that focus exclusively on the documentation of experiments in specific disciplines, such as neurophysiology (odMLtables^7^). While these standards differ in their technical specifications to reflect the unique requirements of each omics field, they share a common set of mandatory elements. These include general project information, such as title, description, and contact details as well as comprehensive descriptions of the biological system under investigation, including sample sources, experimental variables, and the relationships between samples and generated data^3,5^. Recognizing the substantial overlap in these core metadata elements across disciplines, the Proteomics Standards Initiative, which developed MIAPE, intentionally focused only on proteomics-specific requirements. For general metadata components, it adopted existing standards such as MIAME, thereby not only promoting consistency and reducing redundancy but also highlighting the need for coordination across omics fields^2^. In response, the Minimum Information for Biological and Biomedical Investigations (MIBBI) project^8^ was launched with the goal of aggregating minimum information checklists from diverse scientific domains into a centralized resource. By unifying these standards, MIBBI fosters collaboration and coherence within the global data standardization community.

A critical component of effective standardization is the use of a shared, controlled vocabulary to ensure consistent terminology across studies and disciplines. This principle is exemplified by the Darwin Core (DwC)^9^, a widely adopted glossary of terms designed to facilitate the exchange of information about biological diversity. To bridge the gap between biodiversity data and omics research, the Task Group for Sustainable DwC-MIxS Interoperability^10^ was established. Its mission is to strengthen collaboration between the biodiversity and omics communities by aligning minimum information standards with the Darwin Core terminology, thereby enhancing data interoperability and enabling more effective cross-domain data sharing and reuse.

The importance of standardized terminology becomes especially evident If you need to organise and store large amounts of metadata. Inconsistent naming conventions, spelling variations, or the use of synonyms can prevent datasets from being retrieved during searches, significantly limiting their reuse potential. This was demonstrated by the Biosamples Database^1^, which undertook a community-driven curation effort to standardize metadata in existing datasets. By normalizing terminology—resolving spelling differences and harmonizing synonyms—the number of search results for the term *‘sample type’* increased by 25% through improved query coverage. However, such post-hoc curation is labor-intensive and resource-demanding. Introduction of data using a consistent structure and a well-defined, standardized vocabulary would be more efficient. Embedding controlled terminology at the earliest stages of recording ensures greater findability, reusability, and long-term value of omics data.

To meet FAIR principles and adhere to established metadata standards, a universal, open, and sustainable file format is required, which is capable of permanently storing any conceivable experimental design along with its associated sample and condition metadata. The design must account for individual samples, biological and technical replicates, a diverse range of categorical and continuous variables representing experimental factors and additional phenotypic traits, hierarchical and nested experimental structures, and technical details specific to the omics technology employed. Consequently, any standardized interview or metadata recording process intended to populate such a format must be carefully structured to capture all necessary information without overwhelming users, balancing completeness with usability.

In many biomedical laboratories, metadata is initially recorded in paper notebooks, spreadsheets, or electronic lab notebooks (ELNs). As data analysis progresses, this information is often copied or re-entered into additional documents, leading to fragmented, versioned records that may diverge over time and become difficult to reconcile. This lack of integration increases the risk of inconsistencies, data loss, and reduced reproducibility. A centralized metadata management system has the potential to mitigate these issues by providing a single, consistent, and traceable source of truth, significantly simplifying data organization and supporting long-term data integrity and reuse.^11^ While centralized platforms developed at universities such as OpenBIS^12^, Yoda^13^, OMERO^14^ or Coscine^15^ offer powerful solutions for data storage, version control, and querying, their deployment typically requires substantial technical expertise, dedicated IT infrastructure (e.g., databases, web servers), and continuous maintenance. Commercial services, such as OpenProject^16^, that provide hosted IT infrastructure often incur significant costs, thereby restricting access to these solutions for many individuals and institutions. In addition to database-driven systems, file-based approaches have been developed to enable lightweight, accessible metadata storage. One widely used example is MAGE-TAB, a standard for transcriptomic data that consists of two tab-delimited files: the Investigative Description Format (IDF), which captures project-level metadata, and the Sample and Data Relationship Format (SDRF), which links samples to their corresponding data files^17^. Another example is ISA-Tab^18^ tabular format, which was adopted by Fair Data Cube (FDCube)^19^, a technological framework for analyzing multi-omics data. Other approaches involve the development of custom metadata languages, such as MEDFORD and the open metadata Markup Language (odML) framework, which are based on domain-specific markup languages and provide templates to guide users in creating structured metadata^7,20^. Although these file-based solutions often include validators and parsers to detect semantic and structural errors, they generally do not offer features that assist with the actual creation and editing of metadata files such as an input mask or a query. As a result, users must manually construct and maintain these files, which can be error-prone and time-consuming, particularly for researchers without formal training in data modeling or metadata standards^17,20^. This problem is partially mitigated by some third-party tools such as lesSDRF^21^, which provides an interface for creating SDRF files.

In order to overcome the obstacles mentioned above, we here introduce the **F**ai**R E**xperimental **D**esigns (FRED) toolkit for FAIR metadata handling. FRED is based on a machine-readable metadata file format to be stored with experimental data to fully describe an experiment. The toolkit offers a suite of functionalities designed to support the implementation of FAIR data principles. It enables the structured creation of standardized metadata files in YAML format, which are both human- and machine-readable, thereby improving interoperability across diverse systems. Data is semantically validated to ensure metadata consistency and enhance quality, supporting long-term reusability. The toolkit supports cross-file metadata search, addressing the findability requirement by facilitating efficient discovery of relevant datasets. Finally, an application programming interface (API) is provided to enable seamless integration with external systems, including web-based interfaces, thus promoting accessibility and extensibility within broader bioinformatics workflows. The file format and command line tool are supported by descriptive metadata structures and a variable whitelist via a central repository that constitute the management backend for an individual instance of FRED. This architecture is designed to be accessible to non-IT users and requires minimal bioinformatics expertise, allowing for integration with any existing data storage structure.

## Results

### Structure, workflow, and features of the FRED toolkit

FRED presents a practical and user-centric solution for the structured generation and management of metadata in omics research. It is implemented in Python and freely available on GitHub at https://github.com/loosolab/FRED or via the PyPI package index. FRED employs a generic, modular structure (described in the Central Management section) to define and control supported metadata keys, data types, and validation rules. By default, it supports experimental designs involving *n* distinct conditions defined by experimental factors of interest, such as cell type, genotype, disease stage, enrichment, sex, injury or tissue. Individual samples are assigned to these conditions and may represent biological or technical replicates. Each sample can include additional individual or grouped metadata (e.g., age, sex, comorbidities) alongside experimental factors monitored as covariates. During runtime, FRED dynamically accesses a centralized repository of vocabulary (see Central Management section below) to **populate interactive dialogs** and **validate input values in real time**, ensuring metadata consistency and avoiding errors due to user inputs.

The **workflow of FRED** is characterised by the following sequence of events (fig1a): a user describes an experimental design, this description is recorded in a YAML-formatted file, the file is validated during runtime, and the file collection is stored as a metadata repository locally.

These human- and machine-readable YAML files (fig1b) can be integrated seamlessly into existing RDM structures. This is also true for a wide range of downstream tools and analysis pipelines, since most programming languages offer libraries for reading, writing, parsing, and visualizing YAML files. For example, we integrated FRED into the Cardio Pulmonary Institute (CPI) repository (https://bioinformatics.mpi-bn.mpg.de/bcu-repository-bn), a RDM structure intended to store all omics experiments generated within the three sites of the excellence cluster project funded by the Deutsche Forschungsgesellschaft (DFG), and harboring >1000 datasets and metadata files.

**Figure 1.**
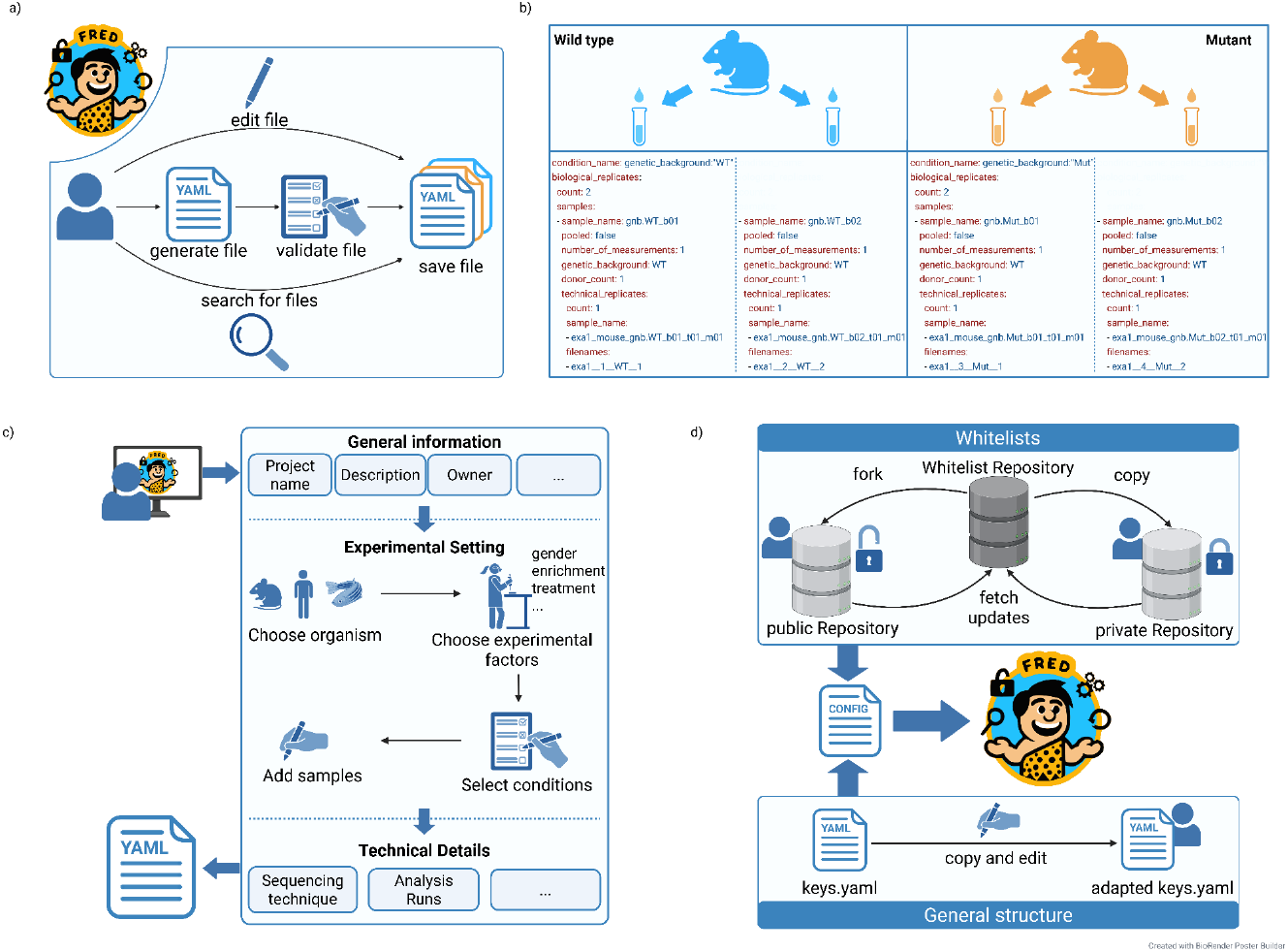
a) The FRED API supports recording, editing, and querying of the metadata stack by a managed vocabulary. b) Exemplary structure of a metadata YAML file for an experiment with two conditions. c) Workflow for metadata recording, supported by FRED via a command line or web interface. FRED’s metadata structure is thematically divided into general information about the project, experimental design and technical details. d) FRED relies on a whitelist repository and the metadata structure YAML (keys.yaml) to facilitate universal adaptability. Forking or copying the whitelist repository (private or public) and editing of the structure YAML enables adaptation of terms and keys.

To support flexible, universal, and hierarchical metadata collection, the FRED toolkit provides a guided, incremental dialog for metadata generation via a standalone web browser (see section below) or command line (fig1c). By guiding users through a structured, intuitive workflow, FRED reduces the risk of missing critical information, promotes consistency across projects, and supports the generation of high-quality, FAIR-compliant metadata from the very beginning of a research study. In addition, FRED automatically generates unique, informative sample IDs and filenames using a standardized format that incorporates project, sample, and experimental information, ensuring each sample is uniquely identifiable and traceable. A summary of all FRED features and a comparison of these features with other approaches and solutions developed in the field is shown in Table 1.

**Table 1:** Comparison of features of metadata tools in various fields.

| Feature/ Tool | OMERO Metadata | Yoda | Coscine | OpenBis | OpenProject | lesSDRF | MARMoSET | FDCube | FRED |
| --- | --- | --- | --- | --- | --- | --- | --- | --- | --- |
| **open source** | yes | yes | yes | yes | free community edition, paid enterprise version | yes | yes | yes | yes |
| **metadata definition** | self defined key-value pairs or tags | pre-defined templates, configurable after consultation with instance data manager | pre-defined templates, configurable after review by the FDM team | defined by instance admin | defined by instance admin | pre-defined structure, not configurable | no fixed structure | ISA metadata schema | pre-defined metadata structure, configurable |
| **validation** | no validation | validation for defined templates, configurable after consultation with instance data manager | validation for defined templates, configurable after review by the FDM team | defined by instance admin | defined by instance admin | validation for defined structure, not configurable | no validation | validation for structure and values | validation for structure and values, configurable |
| **graphical abstract of experimental design** | no | no | no | no | no | no | no | no | yes |
| **containerized** | yes, for development | yes, for development | no | yes | yes | yes, for development | no | yes | yes |
|  | environment | environment |  |  |  | environment |  |  |  |
| **hierarchy** | no hierarchies | low level hierarchies | no hierarchies | hierarchical structure | hierarchical structure | no hierarchies | no hierarchies | hierarchical structure | hierarchical structure, dependencies between keys can be defined |
| **metadata format** | database | json objects | RDF format | database | database | SDRF file | JSON file | JSON file, ISA-Tab file | YAML file |
| **accessibility** | Web interface, commandline, API for clients | Web Interface, API | Web Interface, API | Web Interface | Web Interface | Web Interface | commandline | commandline, API | Web interface, commandline, API |
| **client/server based** | server based | server based | server based | server based | server based | client based | client based | client based | client based |
| **data sharing** | only within OMERO | HTML, OAI-PMH server, json file | RDF file | export metadata as PDF or XLSX | share with registered users | SDRF file | JSON file | JSON file, ISA-Tab file | YAML file |
| **Audience** | facility | facility | facility | facility | facility | single user | single user | single user | single user, workgroups, facility |

### Centralized management of valid designs and vocabulary

The management of metadata structure and whitelists are realized by two code repositories for online/offline/forked usage (fig1d).

All metadata configurations in FRED are centrally managed through a flexible, hierarchical system based on YAML-formatted files. This structure ensures consistency, transparency, and ease of customization across projects and organizations. At the core of the system is the key configuration file ‘keys.yaml’, which defines all permissible metadata keys, their hierarchical placement via indentation, and their structural relationships within the metadata schema.

The metadata schema is thematically organized into three main sections, each represented by a top-level key.

- **project**: Contains general project-level information, including a unique identifier, title, description, and contact details. Contact roles include the data owner, data analyst (if applicable), hosting group, and hosting institution.
- **experimental_settings**: Describes the scientific objectives and structure of the study, specifying how conditions and samples are defined and related. This section allows for multiple experimental setups within one project, enabling the combination of data from different experiments or species.
- **technical_details**: Captures methodological details such as omics technologies used, processing steps, and other technical aspects, ensuring full traceability and reproducibility.

Each key in the key configuration file is associated with a set of properties that define its behavior within FRED. These properties (summarized in Table 2) include data type (e.g., string, list, integer), whether the field is mandatory or optional, allowed value ranges, default values, and validation rules. These settings ensure consistency, prevent invalid entries, and support automated validation during metadata generation.

**Table 2:**
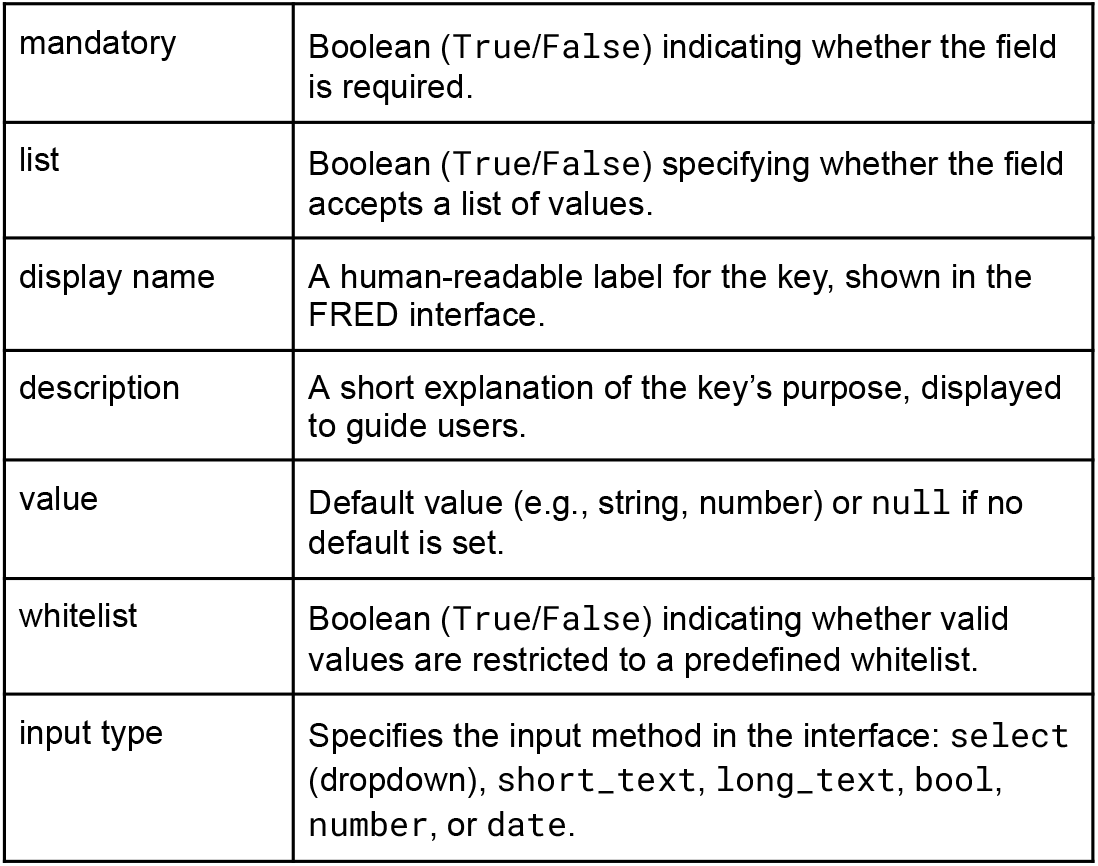
Metadata Key Properties in FRED. This structured definition of metadata properties ensures consistent behavior across all FRED projects and enables automated validation.

|  |  |
| --- | --- |
| mandatory | Boolean (True/False) indicating whether the field is required. |
| list | Boolean (True/False) specifying whether the field accepts a list of values. |
| display name | A human-readable label for the key, shown in the FRED interface. |
| description | A short explanation of the key's purpose, displayed to guide users. |
| value | Default value (e.g., string, number) or null if no default is set. |
| whitelist | Boolean (True/False) indicating whether valid values are restricted to a predefined whitelist. |
| input type | Specifies the input method in the interface: select (dropdown), short_text, long_text, bool, number, or date. |

FRED employs **whitelists** to enforce consistent and standardized user input. Whitelists are stored in a dedicated, publicly accessible repository at https://github.com/loosolab/FRED_whitelists (fig1d). These whitelists are used to restrict input values for string-type fields by providing a predefined set of valid options, thereby preventing inconsistencies due to spelling variations, synonyms, or informal terminology—common causes of failed searches or data misinterpretation.

Users can fork or clone the official whitelist repository to create custom versions or extend existing ones. To maintain alignment with updates from the official repository, GitHub Actions are provided that automate the synchronization process. Users can activate these workflows and, if necessary, adapt them to their specific needs ensuring their local whitelists remain up to date with minimal effort.

Whitelists are categorized into several types to support diverse use cases.

- **plain**: A simple list of valid values for a given key (e.g., a list of tissue types).
- **group**: A dictionary-based structure that organizes longer whitelists into logical categories, improving readability and usability (e.g., grouping diseases by affected organ).
- **depend**: Enables conditional whitelists based on another input. For example, a list of valid gene names depends on the organism (e.g., *Homo sapiens* vs. *Mus musculus*), allowing context-aware validation.
- **abbrev**: Defines abbreviations for values from other whitelists. These abbreviations are used to generate unique, standardized sample file names improving traceability and reducing naming errors.

FRED regularly performs a validation step on each metadata file e.g. triggered by a search metadata operation (see below). The validation process itself consists of two phases (fig 2a), the **Syntactic Validation and the Logical Validation**. The former ensures that each metadata file conforms to the defined structural schema, including correct hierarchical position of metadata keys within the key configuration file structure, presence of mandatory keys, valid data type, and presence in corresponding whitelist. In case of syntactic errors, a message is generated listing all deviations from the schema. Files that fail syntactic validation are excluded from the search to prevent incorrect or misleading results.

**Figure 2.**
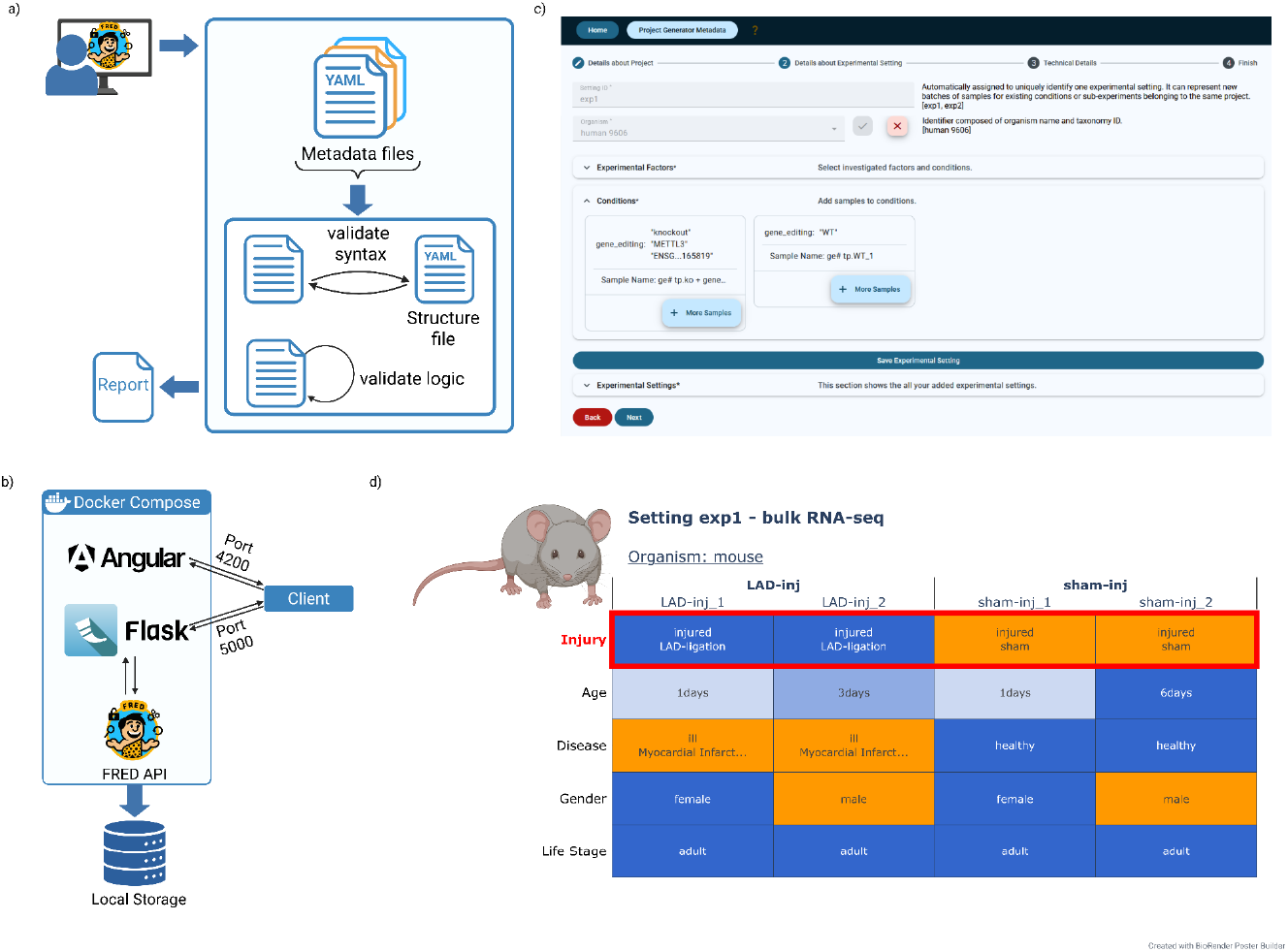
a) FRED provides syntactical and logical validation of metadata files. b) The graphical user interface of FRED consists of two containers, a web app using Angular and a REST-application programming interface using Flask. The containers can be built and started using docker compose. c) FRED provides an interface for external calls, e.g. via a web frontend. d) FRED provides a summarizing visualization of the experimental design with color-coded values. For quantitative or ordered categorical values (e.g. age, time points, concentrations) FRED uses a continuous color gradient to represent the magnitude of differences.The plot includes information about the organism and the technical method used for sample processing (e.g., RNA-Seq, ChIP-Seq) at the top. This is followed by key experimental factors (e.g., genotype, treatment, disease stage) displayed with a red border.

During **Logical Validation**, domain-specific consistency checks are performed to ensure that metadata entries are contextually sound. For example, to verify whether the reference genome specified for a sequencing run is compatible with the organism described in the metadata. Logical validation does not affect the file’s syntax, so it does not prevent the file from being processed. Instead, any inconsistencies trigger a warning message that summarizes all logical issues for the user. These warnings help to identify potential errors or misconfigurations without blocking the search process.

By combining syntactic and logical validation, FRED ensures both structural integrity and semantic coherence of metadata. This dual-layered approach enhances data quality, supports reliable search results, and helps users identify and correct issues early in the data lifecycle—contributing significantly to reproducibility and reusability of omics data.

In summary, the central management system in FRED—combining a structured YAML schema with a modular, extensible whitelist framework—provides a powerful, scalable foundation for standardized metadata collection, iterative structure extension and data integrity. It empowers research teams to maintain high-quality, interoperable data while minimizing errors and supporting long-term data sustainability.

### Guided metadata generation, visualization and search via webinterface

FRED is designed to communicate with the user in two ways. The first method is a command line interface that can be executed directly on a terminal console. This can be used to generate metadata files via a guided dialogue interface or to search for or modify existing files. The second method uses the FRED API and is optimized for users lacking prior experience in computer science or command-line interfaces. It allows researchers to interact with FRED through a graphical user interface implemented using Angular^22^ and provided as a Docker^23^ container (fig2b). The interface guides users through the metadata generation (fig2c), including core FRED functionalities, like real-time validation, visualization of experimental designs, and export of metadata in YAML format. The standalone web frontend is available at https://github.com/loosolab/FRED_standalone.

Metadata files can be stored in a central directory and queried via FRED search functions. This approach is designed to enable efficient discovery and retrieval of metadata across multiple projects and experiments. The search requires two mandatory inputs: a path to a folder to be recursively searched (root directory) and a search string specifying the values to be found in the metadata. The search string can contain one or more values, which are compared against all keys in the metadata files. Users can refine the search by prefixing a value with a specific key name (e.g., organism:”human”) and combining values using logical operators (AND/OR/NOT) with parentheses to control evaluation order.

FRED includes a built-in function to visualize experimental designs as an interactive plot, enabling users to quickly assess structure and composition of their experiments (fig 2d). Each cell is color-coded based on the value it contains, allowing immediate visual identification of similarities and differences across samples. Information can be further collapsed to communicate only the minimal study design, and be exported as an interactive HTML or a PNG file.

In summary, by providing a command line interface as well as self-contained web application, FRED ensures broad accessibility across diverse research teams, including biologists, clinicians, and other non-technical users. Its experimental setup visualization routine is, to the best of our knowledge, unique to the field.

## Discussion & conclusion

FRED presents a practical, scalable, and user-centric solution for structured generation and management of metadata in omics research, fully aligned with the FAIR principles. The toolkit is designed to support researchers across diverse technical backgrounds, offering both a command-line interface for advanced users and a standalone web-based frontend. This dual-access approach ensures broad accessibility, enabling seamless adoption in laboratories with varying levels of technical expertise.

FRED’s metadata structure is comprehensive and modular, encompassing three essential components: general project information, detailed experimental design and sample annotations, and technical metadata. This structure meets essential requirements for omics-domain research^17,24^, while remaining flexible enough to be applied across other types of high-throughput experiments. Unlike many database-driven systems that require expert knowledge to customize or extend, FRED’s hierarchical, YAML-based design permits straightforward adaptation to evolving projects.

Sample tracking is implemented via automated generation of unique, informative IDs derived from experimental factors and project context. These unique IDs are embedded in the metadata, ensuring traceability of samples and derived files across projects.

FRED stores metadata in YAML format to balance machine-readability with human interpretability. This facilitates integration into automated downstream data analysis pipelines or the planned export of metadata to standardized formats like MAGE-TAB (for transcriptomics)^17^ or NCBI Gene Expression Omnibus (GEO)^23^, which will further streamline data submission and promote interoperability across platforms.

In summary, FRED bridges the gap between rigorous metadata standards and real-world usability. By combining a flexible, extensible architecture with intuitive interfaces and strong support for FAIR principles, it empowers researchers to generate high-quality, reusable metadata. With increasing volume and complexity of omics data, computational tools such as FRED play a critical role in ensuring that data are not only generated but also efficiently documented, discoverable, and reusable.

## Methods

### The program FRED

FRED is implemented in Python and hosted on GitHub at https://github.com/loosolab/FRED. It relies on the PyYAML library for reading and writing metadata files and GitPython for managing interactions with the whitelist repository. For the generation of interactive visualizations, FRED uses the plotly Python package. All dependencies are defined in the pyproject.toml file. The tool, along with all required packages, can be installed using pip in a standalone Python environment with no external dependencies. This ensures FRED runs reliably on any standard Linux system without the need for specialized IT infrastructure. A comprehensive user guide is available at https://loosolab.pages.gwdg.de/software/metadata-organizer/, providing clear instructions for installation, configuration, and usage.

### Whitelists

Valid values for metadata fields are stored in a dedicated repository at https://github.com/loosolab/FRED_whitelists in YAML format. To improve performance during runtime, a GitHub Action automatically converts each YAML whitelist into a JSON file. This allows FRED to load and process whitelists more quickly. Users can fork or clone the repository to create custom versions. Pre-configured GitHub Actions enable automatic synchronization with updates from the main repository, ensuring access to the latest standardized terms while maintaining flexibility for local modifications.

### Docker Compose

The standalone version of FRED consists of two containers, a web App using Angular and a REST-API using Flask. The containers can be started with the command ‘docker compose up’. The standalone version is available at https://github.com/loosolab/FRED_standalone.

## Acknowledgement

We would like to thank the staff of the IT group at MPI for heart and lung research, especially Daniel Spothelfer and Vincent Heinz for supporting our virtual computational infrastructure and our Kubernetes cluster.

## Authors contribution

M.L. conceived the project. M.L., C.K. and J.W. designed the software. J.W. implemented the FRED software, N. K. implemented the standalone frontend, P.G. took care of the interoperability with the CPI repository data structure, J.W., C. K. and M.L. wrote the manuscript, M. L. supervised the project.

## Data availability

All code described and respective docker images are available via github as indicated above.

